# Genetic control of Group 3 (K96) capsule synthesis and complement resistance in an extraintestinal pathogenic *Escherichia coli* ST69 strain

**DOI:** 10.1101/2025.09.19.677278

**Authors:** Rachael J. David Prince, Rajesh Bogati, Titan Lytle, Andres Collado Silva, Rachael A. LeBaron, Tiana Sorge, Michael A. Olson, Eric Wilson, David L. Erickson

## Abstract

Extraintestinal pathogenic *Escherichia coli* (ExPEC) often produce capsules belonging to Groups 2 or 3, which contribute to invasive disease in humans and other animals. Group 3 capsule loci contain conserved *kps* genes flanking serotype-specific glycosyltransferase and nucleotide-sugar biosynthesis genes, but little is known about genetic factors that control synthesis, or the specific roles that K96 capsules play in virulence. Previously, we identified a Group 3 serotype K96 capsule in a mastitis-associated strain M12 that is critical for its survival in some host tissues. In this study, we show that the K96 capsule of strain M12 confers resistance to human serum complement. We conducted genetic screens to determine how K96 capsule expression is controlled, which led to two principal findings. First, Group 3 capsule synthesis is unstable. Sequencing of spontaneously appearing mutants revealed potential phase-variable control of capsule expression. Some mutants harbored reversible frameshift mutations in a homopolymeric site within *kpsC*, which were also identified in other K96-encoding ExPEC isolates. Second, a transposon mutagenesis screen revealed that capsule synthesis requires proteins encoded outside of the *kps* locus, including the RfaH antiterminator and OxyR transcriptional regulator, which control activation of an unusually distant promoter. An Δ*oxyR* mutant of strain M12 failed to produce capsule, was extremely sensitive to complement-mediated killing, and avirulent in *Galleria mellonella*.

**Importance:** Group 2 capsules are established extraintestinal pathogenic *Escherichia coli* (ExPEC) virulence factors and are the type most often associated with human isolates. Group 3 capsules have previously been considered a sub-type of Group 2, although nothing was known about factors controlling their expression. Capsule serotype K96-encoding ExPEC strains are increasingly isolated from human infections and inhabit numerous other hosts. It is critical to understand the factors that enable their pathogenic versatility and survival in specific environments. Our study shows that K96 capsule of strain M12 is required for human complement resistance. Additionally, we have identified genetic factors that control Group 3 capsule synthesis, including a potential phase-variable mechanism as well as transcriptional control by OxyR. Co-regulation of Group 3 capsule synthesis genes with genes necessary for oxidative stress resistance may increase the virulence of some versatile ExPEC strains.

## Introduction

Extraintestinal pathogenic *Escherichia coli* (ExPEC) are a leading cause of serious human diseases, including pneumonias and bloodstream infections. Approximately 240,000 yearly deaths are attributable to bloodstream infections due to these bacteria [1]. Many invasive infections begin as a urinary tract infection (UTI), and ∼60% of women experience at least one UTI caused by *E. coli* [2]. ExPEC also cause diseases of livestock and companion animals, and their zoonotic potential is increasingly recognized [3–6]. Expanding resistance to antimicrobial drugs greatly complicates their treatment [7–9]. Individual ExPEC strains may possess a variety of virulence determinants, including metal acquisition systems, adhesins, toxins, and protective structures that shield them from host immune defenses including lipopolysaccharide O-antigen and capsular polysaccharides. However, there are no genetic markers that reliably differentiate ExPEC from commensals or intestinal pathogens [10].

Surface polysaccharides, including K antigen capsules, enhance the virulence of many *E. coli* strains [11]. More than 90 different K antigens are classified into four broad groups based on their mode of assembly and gene arrangement [12, 13]. Group 1 and 4 capsules are polymerized in the periplasm and exported in a Wzx/Wxy-dependent manner, while Group 2 and 3 are polymerized at the cytoplasmic face of the inner membrane and exported via ABC transporters [14]. Specific capsules have diverse functions, including protecting bacteria from serum complement, inhibiting phagocytosis, forming intracellular bacterial communities, resisting antimicrobial peptides, and avoiding detection by macrophage-encoded receptors [15–18].

Most capsules made by ExPEC belong to Groups 2 or 3 [19–23] and previous investigations have focused on the Group 2 capsules. Group 3 *kps* loci contain genes necessary for initiation (*kpsS*, *kpsC*) and export (*kpsD*, *kpsM*, *kpsT*, *kpsE*) of the glycolipids that comprise these capsules, as well as genes for serotype-specific glycosyltransferases that polymerize the sugar subunits. KpsS and KpsC add ketodeoxyoctonic acid to a phospholipid carrier, upon which the capsule is built in the cytoplasm prior to export by the ABC transport apparatus (KpsDMTE). While the synthesis, export, and biological functions of Groups 2 and 3 capsules are presumed to be similar, their genetic organization at the *kps* locus differs. There are two opposing transcriptional units for Group 2 capsules whereas Group 3 capsule genes appear to be arranged in a single operon. Group 2 capsule loci contain *kpsF* and *kpsU* genes that are absent from Group 3, suggesting that they are not needed, or that their functions are fulfilled by genes found elsewhere in the genome. A model for Group 2 capsule expression and regulation has been developed [24], which includes genes encoded outside of the *kps* locus necessary for synthesis [25–27]. Beyond the conserved *kps* genes, nothing is known about genes needed for producing Group 3 capsules or how they are regulated.

We recently showed that *E. coli* strain M12, originally isolated from a clinical case of bovine mastitis, is highly virulent in mouse models of mammary gland infection, ascending urinary tract infection and intraperitoneal sepsis [28]. Strain M12 belongs to the ST69 lineage, like many human ExPEC strains emerging worldwide [29]. This suggests that M12 and perhaps other mastitis-associated strains may be capable of causing ExPEC-like disease in humans.

However, whether strain M12 or other mastitis-associated isolates can resist human innate immune defenses has not been examined. M12 produces a Group 3 capsule which is critical for survival in mouse kidneys during UTI and intraperitoneal infections [28] and moderately enhances fitness in mammary gland infections [30]. This capsule belongs to serotype K96 [31] and likely contains D-glucuronic acid and L-rhamnose [32]. K96 is similar to K54 capsule, produced by the human ExPEC strain CP9, which contains glucuronic acid and rhamnose modified with threonine or serine [33]. K96 and K54 capsules are sufficiently similar that anti-K54 antiserum also binds K96 capsule on strain M12 [28]. The K54 capsule of strain CP9 is important during soft tissue and bloodstream infections, but unlike the M12 capsule, it does not affect survival during UTI in either bladders or kidneys [34]. The K54/96 capsules most commonly occur in strains belonging to the ST69 lineage (like M12), and a recent rapid expansion of human bloodstream isolates that have acquired these Group 3 loci has been documented [35].

While our previous work demonstrated that the unencapsulated M12 Δ*kpsCS* mutant is severely attenuated in some tissues, we did not determine the basis for the virulence defect, and specific functions for K96 capsules more broadly are unknown. Some ExPEC capsules confer protection against serum complement while others do not [36–39], and the effect of capsule on complement-mediated killing likely depends on other factors that vary among ExPEC strains. Although M12 requires capsule to survive in mouse kidneys and the peritoneal cavity, which is consistent with a role in complement resistance, mice are not reliable as a model host to study the role of complement and bacterial resistance factors for human pathogens [40]. In this study we sought to determine whether the K96 capsule affects M12 survival in human blood and serum and to identify genetic and environmental factors that affect Group 3 capsule synthesis.

## Results

### M12 capsule confers resistance to human serum complement

To investigate strain M12 as a potential human pathogen and define the role of its K96 capsule, we first determined its survival in whole human blood. Strain M12 grew very well in blood obtained from two different donors, showing a ∼150% increase over a two-hour period (Figure 1A). The Δ*kpsCS* mutant strain was significantly more sensitive to killing in whole blood compared to the M12 wild-type strain and was not recovered after 60 minutes of incubation (>4 log decrease).

**Figure 1.**
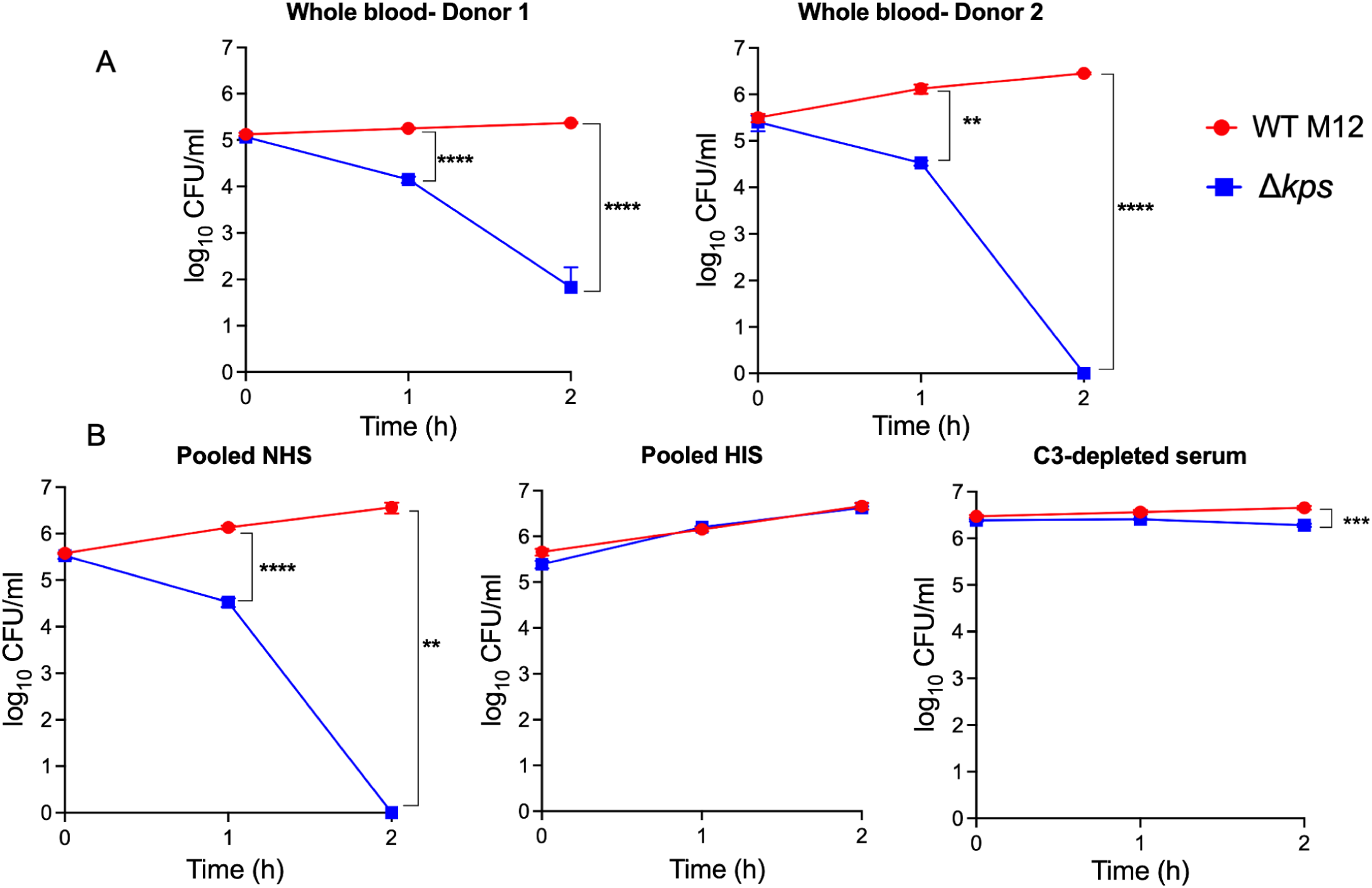
Group 3 K96 capsule confers resistance to human blood and serum complement in strain M12. (A) Survival of M12 wild-type and Δ*kpsCS* mutant in whole human blood obtained from two different donors. (B) Survival of M12 wild-type and Δ*kpsCS* mutant in pooled normal human serum (NHS), heat inactivated serum (HIS) and C3-depleted serum. Asterisks indicate statitistically significant difference from the wild-type strain at that time point, ** = p-value < 0.01, *** = p-value < 0.001 by Tukey’s multiple comparison test.

Next, we compared survival of the wild-type M12 strain with the Δ*kpsCS* mutant in pooled normal human serum (NHS) and in heat-inactivated serum (HIS) (Figure 1B). While the wild-type strain replicated in both NHS or HIS, the Δ*kpsCS* mutant in NHS was rapidly killed. HIS did not kill either strain, suggesting that complement was responsible for death of the mutant strain. We also tested bacterial survival in serum depleted of the complement C3 protein (Figure 1B). Like HIS, C3-depleted serum had very little effect on survival of either the wild-type or Δ*kpsCS* mutant strain, although the wild-type M12 CFU counts were consistently slightly higher than the mutant (Figure 1B).

### Identification of genes required for K96 capsule synthesis

Capsule synthesis can reduce the density of bacterial cells, which allows separation of encapsulated from unencapsulated bacteria using Percoll gradients [41]. As demonstrated in Figure 2A, a gradient consisting of 55% and 75% Percoll separated the encapsulated wild-type M12 strain from the unencapsulated Δ*kpsCS* mutant cells, with the unencapsulated bacteria found in the bottom layer. We used this approach to separate rare, unencapsulated mutants in a previously generated M12 transposon library [30]. In a preliminary screen, single colonies were recovered from the bottom layer of Percoll gradients (putatively containing unencapsulated bacteria). We determined the location of the transposon insertion sites for 26 mutants that failed to produce capsule, which revealed fifteen unique transposon insertion sites (Table 1). These included *kpsD*, *kpsT*, *kpsE*, *kpsS*, *kpsC*, *rmlC*, and putative glycosyltransferase genes within the *kps* locus, suggesting that this approach could be used at a larger scale to identify relevant genes. We also obtained mutants with insertions outside of the *kps* region (Table 1), representing genes whose function has not been associated previously with capsule production.

**Figure 2.**
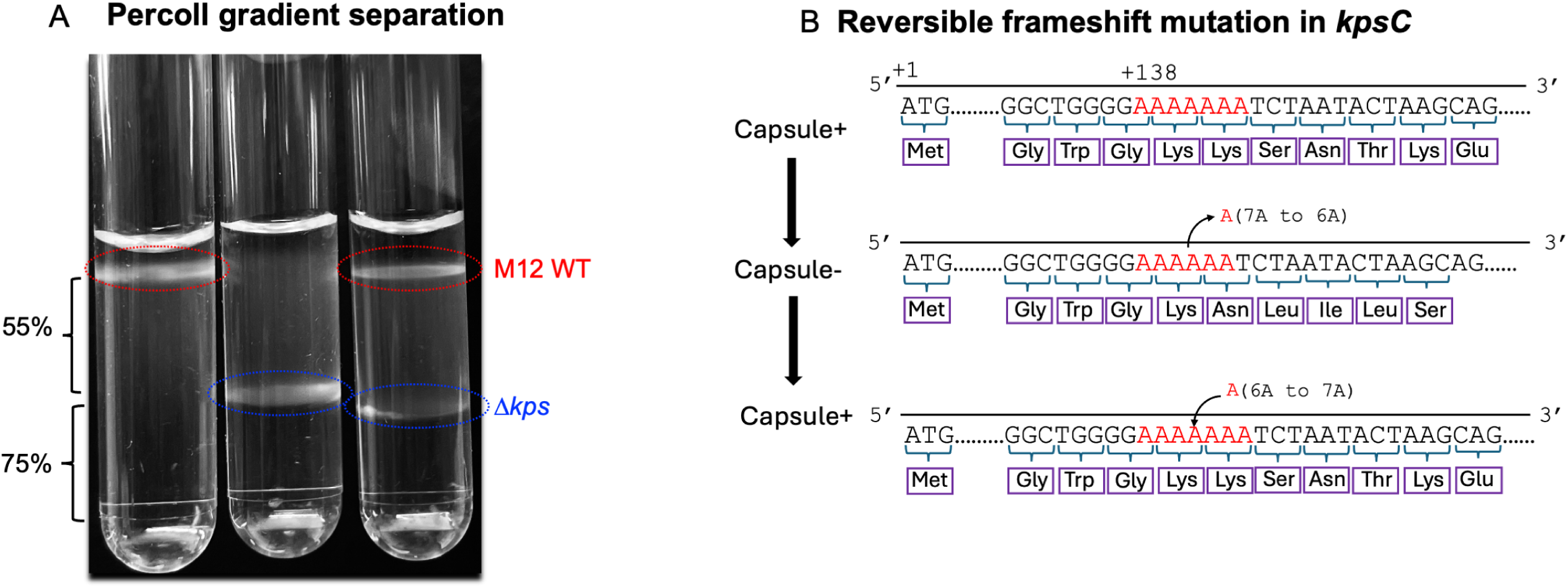
Isolation of mutants defective for K96 capsule synthesis using Percoll gradients. (A) M12 wild-type (left), Δ*kpsCS* mutant (middle) and mixture of both strains (right) were applied to Percoll gradients (55% and 75%) to demonstrate separation of the encapsulated (top layer) from unencapsulated bacteria (bottom layer). Percoll gradients were used to isolate individual transposon mutants that do not produce capsule from an M12 Tn5 library (listed Table 1). (B) Deletion identified within a homopolymeric site of adenosines in *kpsC* in an *mlaC*::Tn5 transposon mutant as well as in two spontaneous mutants derived from M12 wild-type parent strain (See Table 2). Subculturing and Percoll separations of two unencapsulated frameshift mutants both yielded capsule-producing revertants, whose *kpsC* genes were in frame as shown.

**Table 1.** Genes disrupted in unencapsulated Tn5 isolated from Percoll gradients.

| Gene name | Number of unique insertions | Function | Capsule restored by complementation |
| --- | --- | --- | --- |
| *kpsD* | 2 | Capsule biosynthesis/transport protein | Not tested |
| *kpsT* | 1 | Capsule ABC transporter ATP-binding protein | Not tested |
| *kpsE* | 3 | Capsule export inner membrane protein | Not tested |
| *wcaA* | 4 | Putative glycosyltransferase | Not tested |
| *wsaE* | 1 | Putative glycosyltransferase | Not tested |
| *rmlC* | 1 | dTDP-4-dehydrorhamnose 3,5-epimerase | Not tested |
| *kpsC* | 3 | Capsule biosynthesis/initiation protein | Not tested |
| *kpsS* | 2 | Capsule biosynthesis/initiation protein | Not tested |
| *ispA* | 1 | Farnesyl diphosphate synthase | yes |
| *arnC* | 1 | Undecaprenyl-phosphate 4-deoxy-4-formamido-L-arabinose transferase | no |
| *yjhH* | 1 | putative 2-dehydro-3-deoxy-D-pentonate aldolase | yes |
| *mlaC* | 1 | Outer membrane asymmetry protein | no |
| *yahL* | 1 | Hypothetical protein | no |
| *ybfL* | 1 | transposase | no |
| *oleD_1* | 1 | Oxoalkanate reductase | no |
| *rbfX* | 1 | O-antigen transport | no |
| *yahK* | 1 | Aldehyde reductase | no |

**Table 2.** Mutations identified in spontaneous unencapsulated mutants.

| Mutant | Gene | Predicted function | Nature of mutation | Effects |
| --- | --- | --- | --- | --- |
| CM1 | *wsaE* | glycosyltransferase | Single nucleotide substitution | W128* |
| CM3 | *wcaA* | glycosyltransferase | Single nucleotide deletion | frameshift |
| CM12 | *ugd* | UDP-glucuronic acid synthesis | IS629 insertion | gene disruption |
| CM13 | *wsaE* | glycosyltransferase | IS629 insertion | gene disruption |
| CM15a | *wcaA* | glycosyltransferase | Single nucleotide substitution | H125Y |
| CM16a | *rbsR* | ribose operon repressor | Single nucleotide substitution | A241V |
|  | *mreB* | cell shape determining protein | Single nucleotide substitution | P115L |
| CM17A | *wcaA* | glycosyltransferase | Single nucleotide insertion | frameshift |
| CM18B | *kpsC* | capsule initiation | Single nucleotide deletion<sup>1</sup> | frameshift |
| CM19A | *wcaA* | glycosyltransferase | Single nucleotide deletion | frameshift |
| CM20A | *kpsC* | capsule initiation | Single nucleotide insertion | frameshift |
| CM21A | *wsaE* | glycosyltransferase | Single nucleotide substitution | W383C |
| CM22A | *kpsS* | capsule initiation | Single nucleotide substitution | I814F |
| CM23A | *wcaA* | glycosyltransferase | Four nucleotide deletion | frameshift |
| CM24A | *kpsC* | capsule initiation | Single nucleotide substitution | L675* |
|  | *irp1* | non-ribosomal peptide/polyketide synthase HMWP1 | Single nucleotide insertion | frameshift |
| CM25A | *gspC2-k96-11* | Type II secretion and *kps* locus genes | 25,700 nucleotide deletion | gene loss |
| CM26A | *kpsC* | capsule initiation | Single nucleotide deletion<sup>1</sup> | frameshift |

<sup>1</sup>Identical mutations in tandem A repeat region.

We attempted to complement mutants containing non-*kps* Tn5 insertions with functional copies of the disrupted gene on high-copy plasmids. This restored capsule synthesis in the *yjhH*::Tn5 and *ispA*::Tn5 mutants (Supplementary Figure 1). However, other mutants outside of the *kps* locus were not complemented by this approach. This could be due to polar effects of the transposon insertions or non-optimal expression of the relevant genes from the high-copy plasmid. We were particularly interested in the *mlaC* gene because of its role in outer membrane asymmetry. Therefore, we inserted the entire *mlaFEDCB* operon on a low-copy plasmid into the *mlaC*::Tn5 mutant and this also failed to restore capsule synthesis. We generated a new Δ*mlaC* deletion mutant, which produced capsule like the wild-type M12 strain (data not shown), showing that the Mla system is not required for capsule production. We then sequenced the genome of the *mlaC*::Tn5 transposon mutant to determine if it harbored additional mutations that affect capsule synthesis. In addition to the transposon insertion in *mlaC*, we identified a single nucleotide deletion within *kpsC* at a repetitive section of adenosines (7A to 6A), resulting in a frameshift mutation (Figure 2B), explaining the lack of capsule production in the *mlaC*::Tn5 mutant.

Frameshift mutations at homopolymeric sites can facilitate variable expression of capsules in other bacteria [42, 43]. Therefore, we sought to estimate the number of unencapsulated bacteria present in saturated wild-type M12 cultures and investigate the genetic basis for their appearance. When grown in LB, the proportion of the cells that were recovered from the bottom of the gradient was 0.0043±0.0036% of the total bacteria in the tube. In comparison, when the bacteria were grown in LB containing 2% bile salts, which we previously showed causes loss of capsule [28], 12.99±9.17% were found in the bottom layer. We observed that most single colonies recovered from the bottom fraction resumed capsule synthesis upon regrowth, suggesting transient loss of capsule gene expression rather than inactivating mutations. However, we identified 16 distinct mutants that did not produce capsule upon re-growth in LB media.

Sequencing revealed that all but one of these mutants contained a mutation within the *kps* locus, including single nucleotide deletions, substitutions, IS629 insertions, and larger deletions (Table 2). Two independently derived mutants had the same deletion within the homopolymeric site in *kpsC* (Table 2, Figure 2B). To determine if this *kpsC* frameshift mutation is reversible, we added a barcoding plasmid to the mutant to prevent and detect contamination and repeated the Percoll separation, this time recovering and subculturing the top portion of the gradient. After five passages, we recovered a population of capsule-producing revertants derived from the barcoded mutant. Resequencing showed that the homopolymeric site in *kpsC* had been restored (Fig 2B). Collectively, this suggests that Group 3 capsule production may be regulated by the regular appearance of inactivating mutations, including via rare slipped-strand mispairing in *kpsC*. To determine whether this occurs naturally, we retrieved all the *E. coli* genome sequences that contain complete K96 loci from NCBI (58 total strains, Supplementary Figure 2A). Most (51) of them are complete, with SNPs present throughout that may or may not impact capsule synthesis positively or negatively. However, five strains were identified that contained an identical single nucleotide deletion within *kpsT*, one strain with an IS element within *k96-05*, and one with single nucleotide insertions/deletions in *wcaA*, *wsaE*, and *kpsC*. (Supplementary Figure 2). Thus, at least 12% of these isolates likely do not produce K96 capsule. Notably, two strains show the identical 7A to 6A frameshift mutation in *kpsC* that we obtained in our experiments with M12 (Supplementary Figure 2).

To identify additional genes required for capsule synthesis, we used Percoll separations to enrich for unencapsulated mutants present in our transposon mutant library, followed by high-throughput sequencing of the transposon junctions. The insertion sites were mapped to the M12 genome and genes overrepresented in the unencapsulated fraction relative to the entire library are listed in Supplementary Table 1. According to this analysis, all genes in the *kps* locus except one predicted glycosyltransferase (*k96-09*) and *rmlB* appear to be required for capsule synthesis (Figure 3A). Putative capsule-associated genes outside of the *kps* locus include several involved in carbohydrate transport and metabolism, hydrogenases, and transcriptional regulators such as OxyR and RfaH (Supplementary Table 1).

**Figure 3.**
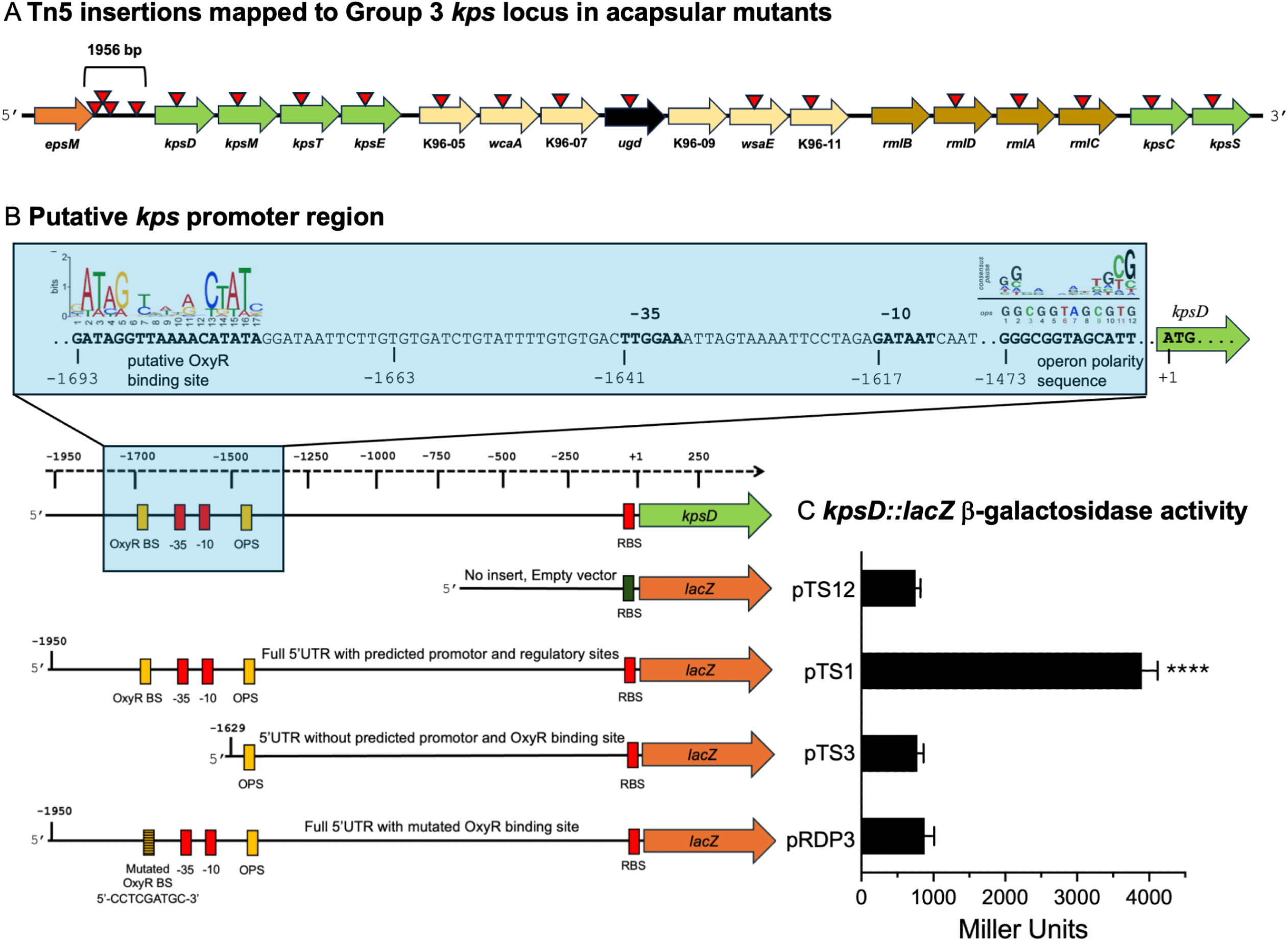
Genes within the *kps* locus required for capsule synthesis including potential *kpsD* promoter region. (A) Structure of the *kps* locus, including those that were identified in the TnSeq analysis as necessary for capsule production indicated with triangles. Many insertions in the unencapsulated transposon mutant population mapped to the intergenic region between *kpsD* and the type II secretion (T2SS) gene *epsM*. (B) Promoter features identified within the T2SS-*kpsD* intergenic region, including putative −35/−10 regions ∼1600 bp upstream of *kpsD*. An operon polarity sequence (ops) necessary for RfaH-mediated transcription of long operons is located 1473 bp upstream of *kpsD*. The consensus ops element from xx is included for comparison. A putative OxyR binding site 5’ to the −35/−10 sequence was identified by comparison with the consensus binding sequence from [58]. (C) *kpsD::lacZ* β-galactosidase assays demonstrate that the predicted promoter region and 5’ UTR including the OxyR binding site activate transcription (pTS1), whereas a fragment that omits the putative −35/−10 region (pTS3) does not. Mutation of the OxyR binding site (pRDP3) results in activity equal to the empty vector control plasmid (pTS12). **** = p-value < 0.0001 by Student T-test compared with pTS12.

Mapping of the transposon sequencing reads to the M12 genome revealed that many of the transposon insertions in the capsule-negative population occurred within the 1956 nucleotide region between *kpsD* and type II secretion genes (Figure 3A). There is a *kpsM*’ pseudogene fragment that is not predicted to yield a protein, as well as a hypothetical open reading frame just upstream of *kpsD* within this region (Supplementary Figure 2). We deleted this region (−1450 to – 50 relative to *kpsD*) from the chromosome of strain M12. This mutant produced capsule (data not shown), demonstrating that neither the *kpsM*’ fragment or the hypothetical protein are required. We also located potential –35 and –10 promoter sigma factor binding sites 1641 and 1617 nucleotides from the KpsD start codon (Figure 3B). There is an operon polarity sequence (ops) necessary for RfaH-mediated transcription elongation [44] downstream of the predicted –35 and – 10 promoter site. We also identified a putative OxyR binding site 5’ to the predicted –35 and –10 promoter site (Figure 3B).

To determine whether this region could activate transcription, we created a reporter plasmid with the 1950 nucleotide *kpsD* upstream region fused to the *lacZ* gene. This fragment was sufficient to induce β-galactosidase activity in strain DH5α, which lacks native β-galactosidase. However, when the putative promoter and OxyR binding sites were omitted from the reporter plasmid, β-galactosidase activity decreased to levels similar to the plasmid-only control (Figure 3C). We also created a plasmid that retained the promoter, but the putative OxyR binding site was altered, which also resulted in β-galactosidase expression indistinguishable from the plasmid-only control (Figure 3C). These results suggest an unusually long 5’ untranslated region in the *kps* operon and that OxyR may be required for activating Group 3 capsule gene expression in strain M12.

### RfaH and OxyR are required for Group 3 Capsule Production

The TnSeq results and the presence of an ops element downstream of the potential transcriptional start site suggested that RfaH may be required for Group 3 capsule synthesis. We confirmed this hypothesis by creating an Δ*rfaH* mutant strain, which failed to produce capsule. Capsule production was restored in the mutant complemented with the functional *rfaH* gene on a plasmid (Figure 4A). TnSeq and reporter assays also indicated a requirement for OxyR to produce K96 capsule. We deleted *oxyR* from strain M12 and this mutant strain also failed to produce capsule (Figure 4A). The complemented Δ*oxyR* mutant strain produced capsule, although at lower levels than the wild-type strain (Figure 4A).

**Figure 4.**
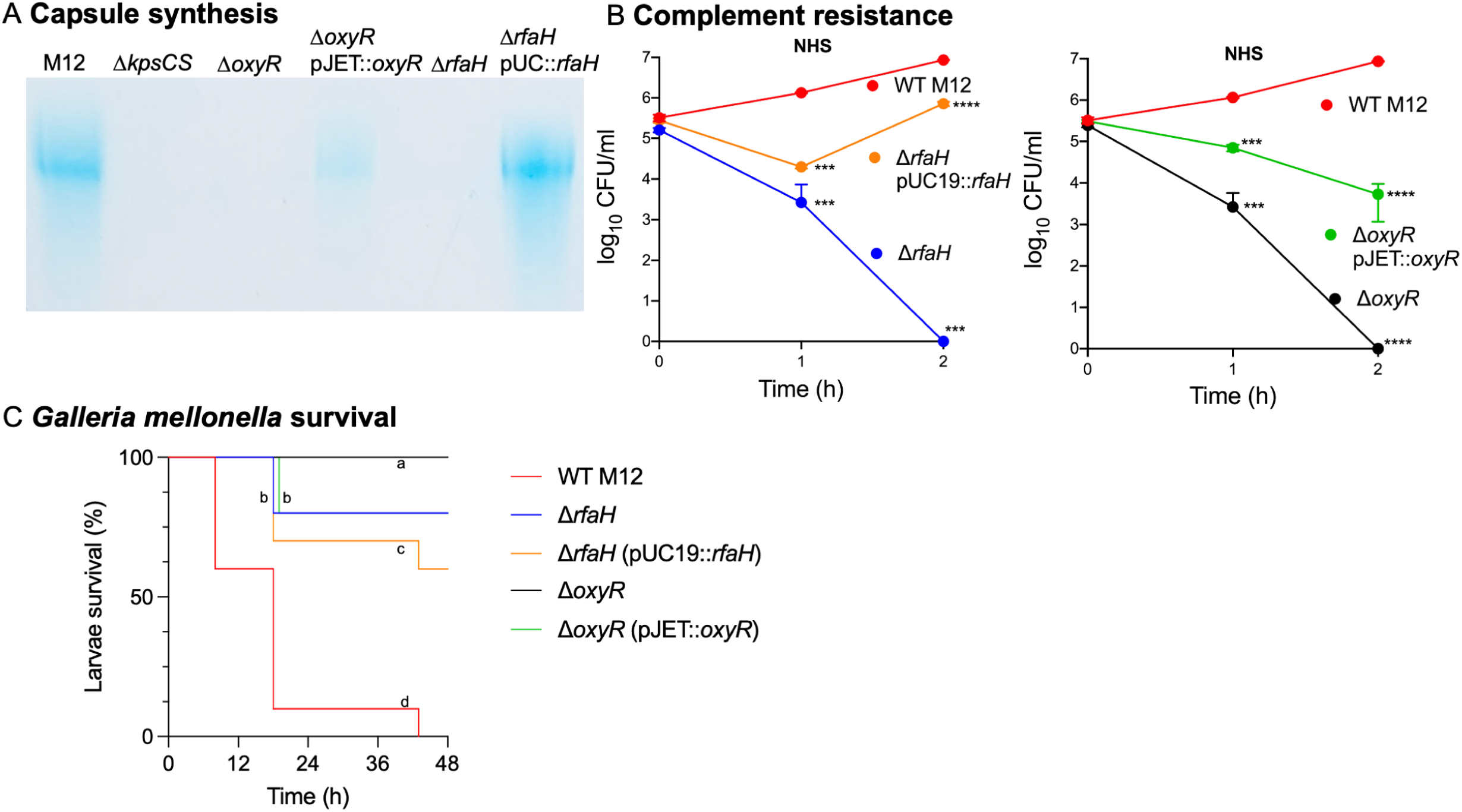
RfaH and OxyR are required for Group 3 K96 capsule production, serum survival, and virulence in *G. mellonella*. (A) Capsule production in wild-type M12, Δ*kpsCS,* Δ*rfaH,* Δ*oxyR,* and complemented mutants detected by SDS-PAGE and Alcian blue staining. (B) Bacterial survival of the same strains incubated in NHS. Asterisks indicate a statistically significant difference from the wild-type strain at the given time by Tukey’s multiple comparison test *** = p-value < 0.001, **** = p-value < 0.0001 (C) Survival of *G. mellonella* larvae infected with 10^4^ CFU of the same strains. Different letters indicate significant differences between strains (log-rank test, P<0.05 after Sidak’s correction).

Since capsule is required for complement resistance we tested whether the Δ*rfaH* and Δ*oxyR* mutant strains survive in NHS. Consistent with their role in capsule synthesis, both mutants were very sensitive to NHS but not HIS (Figure 4B). The complemented *oxyR* mutant strain had an intermediate sensitivity to complement, consistent with incomplete restoration of capsule synthesis. Capsule is also required for virulence of strain M12 in *Galleria mellonella* infections [28]. When compared to the wild-type M12, both the Δ*rfaH* and Δ*oxyR* mutant strains were extremely attenuated in this infection model (Figure 4C).

## Discussion

In this study, we demonstrated that strain M12 is resistant to human serum complement if it expresses its K96 capsule (Figure 1). Although it was isolated from a case of bovine mastitis, these findings and previously reported experiments in animal models of UTI and sepsis [28], strongly suggest that it can cause invasive disease in humans or other animals. Strains whose genomes are extremely similar to M12 have been isolated from a variety of human infections and other environments, further supporting this claim (Supplementary Table 2). Strains that cause bovine mastitis have not previously been considered potential human pathogens. Although rarely possessing genes encoding Group 2 or 3 capsules, mastitis-associated *E. coli* strains are often highly resistant to bovine serum [45], which likely increases the severity of mammary gland infections and invasive disease in cattle. Further work is needed to understand the mechanisms controlling complement resistance in this group of bacteria. Although only 10-20% of mastitis-associated strains belong to phylogroups B2 or D, they often carry virulence genes contributing to adherence, iron acquisition, and immune evasion and carry out an ascending infectious process similar to UTI [46]. Their relevance as sources of human disease will likely increase in tandem with the unfortunate popularization of unpasteurized milk and other dairy products.

ExPEC capsules play important roles in evading host defenses [15, 17, 18, 47]. Genetic variation within the *kps* gene clusters is substantial and some K types are strongly associated with invasiveness [35]. These variations emerge due to horizontal gene transfer and recombination [48]. K96 capsules are associated with high invasiveness among bloodstream ExPEC isolates [35], although the molecular basis is unknown. The precise mechanism whereby K96 capsule confers serum resistance remains to be determined. One strong possibility is that it recruits the control protein Factor H, which inactivates spontaneously generated C3b at the cell surface [49] to prevent activation of the alternative pathway. The glucuronic acid in the K96 capsule confers a polyanionic structure and therefore might recruit Factor H [50], similar to the purported function of sialic acid-containing capsules such as K1 [51]. It is also possible that the K96 capsule does not promote C3 inactivation but rather sequesters the membrane attack complex far from its target. In this case, the deposition of complement proteins could still opsonize the bacteria leading to enhanced phagocytic killing. K96 capsule could also shield antibody targets important to activation of the classical pathway and opsonization. Further experiments are required to distinguish between these possibilities.

The extreme sensitivity of the M12 Δ*kpsCS* mutant to complement was somewhat unexpected as M12 also contains the gene cluster encoding O-antigen belonging to serotype O15. This O-antigen is sufficient to confer significant resistance to bovine serum in mastitis clinical isolate P4 [52] but apparently not in strain M12. It is possible that strain M12 produces only rough LPS, or that the disparate outcomes are related differences between bovine and human complement.

There are certain environments wherein maintaining at least a fraction of unencapsulated cells would be beneficial, such as in the presence of some phages or capsule-targeting antibodies. We have previously shown that growth of encapsulated M12 bacteria in the presence of bile salts is delayed, likely because of insufficient metal uptake [28]. Downregulation or inactivation of capsule synthesis could also potentially improve attachment to specific receptors or surfaces.

Mechanisms that ensure capsule phenotypic diversity would therefore be expected. We readily isolated spontaneous unencapsulated M12 mutants with a variety of changes within the *kps* locus (Table 2). This included more than one isolate with a frameshift mutation in a homopolymeric site in *kpsC*, which we showed was reversible (Figure 2). This finding underscores the caution that should be taken in interpreting transposon mutant phenotypes, as demonstrated by the *mlaC*::Tn5 mutant that also had a *kpsC* frameshift mutation. Several of the mutants we isolated (Table 2) as well as naturally occurring strains (Supplementary Figure 1) have IS elements disrupting the *kps* operon. Similar IS-mediated capsule inactivation has been noted for *Klebsiella pneumoniae* strains in gut environments [53, 54]. These mutants may be better adapted to survival in intestines but would be eliminated in some extraintestinal sites. Future studies should address whether Group 2 or 3 capsule genes in ExPEC strains more broadly are frequently inactivated or undergo phase variation and whether fitness of capsule-negative M12 strains is altered during intestinal colonization. Wider sampling from diverse sources could determine whether the other specific mutations we identified in M12 (Table 2) arise in other strains.

Of the 17 genes within the M12 K96 locus (Figure 3), 15 were predicted by TnSeq to be required for capsule production, including all of the conserved Group 3 *kps* genes. One gene that appeared not to be required was *k96-09*, encoding a probable glycosyltransferase. There are six predicted glycosyltransferases, the precise functions of which are not known. The initial transposon library had few insertions in the *k96-09* gene [30], potentially masking its importance. The *rmlB*, which encodes dTDP-glucose 4,6-dehydratase, also appears not to be required. To produce K96 antigen, UDP-glucose dehydrogenase (*ugd*) must generate UDP-glucuronic acid, and dTDP-rhamnose must be produced by the *rmlBDAC* gene products. This gene contained many insertions in both the original library as well as the unencapsulated fraction. The same enzyme is also encoded within the *wec* locus in the M12 genome for producing enterobacterial common antigen [55]. This redundant gene could be sufficient for capsule synthesis.

Our study also identified other capsule-related genes beyond the *kps* locus. We confirmed *ispA* to be necessary for capsule synthesis (Table 1, Supplementary Figure 1). IspA synthesizes farnesyl diphosphate, which is a precursor to undecaprenyl phosphate [56]. Undecaprenyl phosphate is important for building of peptidoglycan, lipopolysaccharide, and some types of capsule polysaccharides [57]. However, Group 2 and 3 capsules are assembled in the cytoplasm on a phosphatidylglycerol lipid anchor [13]. It is possible that the ABC transport apparatus needed to produce K96 antigen is unable to function without *ispA*. The 2-dehydro-3-deoxy-D-pentonate aldolase YjhH is also necessary. Neither of these genes were detected in the TnSeq analysis, suggesting that our screen was not saturating and additional genes could be identified.

Similar to Group 2 capsules [25], we showed that the transcriptional antiterminator RfaH is required for Group 3 capsule synthesis in strain M12. More surprisingly, our study revealed that *oxyR* is required for Group 3 capsule production, which has not been previously reported for any *E. coli* capsules. OxyR belongs to the LysR family of DNA-binding transcriptional regulators and responds to reactive oxygen species. These oxidize specific cysteine residues, resulting in formation of a tetramer of OxyR subunits, and binding to a conserved DNA element. This binding activates oxidative defence genes like *katG*, *aphCF*, *oxyS*, *gorA*, and *dps* [58] to neutralize the threat. OxyR-mediated Group 3 capsule regulation is reminiscent of typhoid Vi capsule [59], which is also assembled in the cytoplasm and exported by an ABC-transport system [13], and in which polymorphisms arise frequently [60]. OxyR also activates other virulence-associated factors that do not have obvious oxidative-stress resistance roles, including Ucl fimbriae carried by some ExPEC [61]. Our experiments showed that an Δ*oxyR* mutant was unable to produce capsule (Figure 4). Furthermore, when the putative OxyR binding site was altered, the reporter plasmid containing the *kpsD* promoter yielded less β-galactosidase activity (Figure 3C). While this is suggestive of direct OxyR binding and transcriptional activation, this remains to be definitively shown. Regardless, OxyR-mediated direct or indirect activation may be a key feature that distinguishes Group 2 from Group 3 capsules. The ability to activate capsule synthesis alongside oxidative stress resistance genes may enhance resistance of M12 and other Group 3 capsule-producing strains to phagocytes.

In summary, the K96 capsule confers high levels of resistance to human complement but is likely detrimental in other environments, which may select for frequent capsule gene inactivation. In the context of extraintestinal infections, Group 3 K96 capsule genes does not dramatically affect M12 colonization of bladders or mammary glands [28, 30]. However, initial colonization of the glands results in a strong influx of neutrophils to the lumen, which play critical roles in limiting infection [62, 63]. In instances where neutrophils are unable to kill all the bacteria in these environments, the reactive oxygen species they release may trigger more capsule production, which could increase the likelihood of further invasion and systemic spread of Group 3 capsule-producing ExPEC. This may also occur in *G. mellonella* infections, in which hemocytes attempt to kill diverse bacteria in part through oxidation [64–66]. In this model, the M12 *oxyR* mutant is severely attenuated (Figure 4), because it is sensitive to oxidative stress and also unable to activate capsule production.

## Materials and Methods

### Bacterial strains and conditions

M12 wild-type and mutant strains were routinely cultured at 37°C in Luria-Bertani (LB) medium, supplemented when needed with kanamycin (50 µg/ml), chloramphenicol (10 µg/ml) or ampicillin (100 µg/ml). The M12 transposon library was created as previously described [30].

### Blood and serum survival assay

Blood was collected from five healthy donors by venipuncture into serum-separating tubes and was centrifuged at 2000 rpm for 15 minutes. The extracted normal human serum (NHS) was pooled, aliquoted, and stored at –80°C for further use. Heat-inactivated serum (HIS) was prepared by heating the NHS at 56°C for 30 minutes. Whole blood was also obtained from two healthy volunteers by venipuncture into sodium heparin tubes. Cultures to be tested were subcultured, grown to exponential phase, and standardized to 5 x 10^8^ CFU/ml. Serial dilutions were performed in phosphate-buffered saline (PBS) and 5×10^5^ CFU were mixed in triplicate 100 μl volumes of heparanized whole blood or serum. Samples were taken immediately and again at 60 and 120 minutes, followed by serial dilution, plating, and incubation at 37°C overnight to enumerate bacterial survival. The study was approved by the Institutional Review Board of Brigham Young University (IRB2021-375).

### Percoll gradient separation of encapsulated and unencapsulated bacteria

The Percoll gradient separation method was adapted from [41]. Solutions of 75% and 55% Percoll (1 ml each) were layered in polystyrene culture tubes. Then, 200 μl culture volumes (from the M12 transposon mutant library or the wild-type parent M12 strain to isolate spontaneous mutants) were dispensed at the top of the gradient and centrifuged at 3000xg for 30 minutes at 6°C using swing-bucket rotor. To recover unencapsulated populations from the bottom 75% gradient, the tube was sterilized with 70% ethanol and punctured with sterile tweezers. The bottom fraction of the gradient was collected, diluted, and spread on LB plates to recover individual colonies. Individual transposon mutant colonies were screened for loss of capsule synthesis by alcian blue staining (see below) or by Percoll gradient separation. The transposon insertion sites for individual mutant colonies was determined by arbitrary PCR and Sanger sequencing (Eton Biosciences) as previously described [67].

To isolate spontaneous mutants, the bottom fraction from a Percoll separation was subcultured directly into fresh LB broth, grown to saturation, and the gradient separation was repeated before serial dilution and plating. Single colonies were screened for loss of capsule synthesis by Alcian blue staining. DNA from one unencapsulated colony from each experiment was isolated for whole bacteria genome sequencing. Illumina sequencing was performed (SeqCenter), and in some instances long-read Nanopore sequencing (Plasmidsaurus) to identify larger insertions or rearrangements. Polymorphisms were identified by mapping the Illumina reads to the complete M12 genome using Mapping and Find variations/SNP functions within Geneious software platform (Biomatters). Mutants with frameshift mutations in *kpsC* were tested for their ability to revert to capsule synthesis by inserting a barcoding plasmid [68] encoding chloramphenicol resistance to prevent contamination. The barcoded mutants were sequentially grown to saturation in LB broth containing chloramphenicol, followed by Percoll separation and recovering the top portion. After four passages, the top portion was diluted and plated, and single colonies were tested for capsule production. Whole-genome Illumina sequencing of putative revertants was performed by SeqCenter. The frameshift mutants and the putative revertants were also separately verified by PCR amplification and Sanger sequencing (Eton Biosciences).

### Tn-seq analysis

Three aliquots of the library were incubated for 2 h in LB and applied to Percoll gradients to separate the unencapsulated subpopulations, which were recovered and grown for 8 h before harvesting the DNA. The same library was grown in LB but without Percoll separation as the control samples. Genomic DNA extraction, fragmentation, and addition of Illumina adapters with barcodes were performed as described previously [30]. Illumina 100 bp reads were generated on a NextSeq 2000 P1 Illumina reads (single ends) at SeqCenter. The sequences corresponding to each sample were separated using their barcodes, and mapped to the M12 genome using the STAR aligner [69]. Raw count tables for each gene were analyzed using the DESeq2 package in Geneious (Biomatters). The fold change values were log_2_ transformed, and *P* values calculated using a negative binomial distribution performed by DESeq2. Candidate genes required for capsule synthesis were identified by a log_2_-fold change >2 and a *P* value < 0.05.

### Capsule isolation and Alcian blue staining

Capsules were released from pelleted cells by heat treatment (55°C for 30 minutes) and analyzed using polyacrylamide gel electrophoresis as previously described [28, 30]. Negatively charged capsule in the gels was visualized by staining with 0.125% Alcian blue in 40% ethanol/5% acetic acid.

### Gene deletion and complementation

Primers used in creating defined mutants and complementation plasmids are listed in Supplementary Table 3. Gene deletions were generated using the pFOK allelic exchange plasmid as described [70]. Briefly, 500 base pair regions flanking the gene to be deleted were amplified and inserted with NEBuilder^®^ HiFi DNA Assembly (NEB) into EcoR1-linearized pFOK. Merodiploids were selected following conjugation with donor strain on LB agar plates containing 50 μg/mL kanamycin. Counterselection was performed at 30°C on no-salt LB plates with 20% sucrose and 0.5 μg/mL anhydrous tetracycline. Potential mutants were identified by colony PCR and verified by Sanger sequencing. Mutants were complemented by cloning the disrupted gene into a multicopy plasmid pJET1.2 (Thermo Scientific) or pUC19. Plasmids were sequence verified and transformed into the relevant mutant strain by electroporation.

### β-Galactosidase assay

Reporter plasmids were created by PCR amplification of the plasmid backbone and inserts followed by Gibson assembly. Plasmid pTS1 contains the entire putative *kpsD* promoter including the ribosome binding site and start codon fused to *lacZ* in pTS12 [71]. pTS3 was created from pTS1using primers that exclude the putative OxyR binding and −35/−10 sites. Site-directed mutagenesis was used to alter the OxyR binding site in pTS1 to create pRDP3. The reporter plasmids were verified by Oxford Nanopore sequencing (Plasmidsaurus) and β-galactosidase levels were measured by Miller assay as described previously [72].

### *Galleria mellonella* infections

*G. mellonella* larvae were purchased from Best Bet, Inc. (Blackduck, MN), stored at 15°C in the dark, and used within 2 weeks. An overnight culture of bacteria was subcultured and grown to an absorbance of 1.0 at 600 nm and diluted in PBS. Larvae without any evidence of melanization were injected through the left hindmost proleg with 10 μl of the inoculum containing 10^5^ CFU using a Hamilton 701RN syringe and a 30-gauge needle. Control larvae were injected with 10 μl of sterile PBS. The larvae were incubated at 37°C, and survival was monitored over a 48-h period.

### Phylogenetic analysis

Genomes similar to M12 were identified using the Similar Genome Finder Service at the Bacterial and Viral Bioinformatics Resource Center. The default settings were used (maximum 50 genomes returned, maximum distance of 0.05) and the list was manually curated to remove duplicate entries. Complete K96 *kps* loci were identified by BLASTN search at National Center for Biotechnology Information using the M12 reference sequence. Sequences with >99.9% identity across the entire region were selected. Multiple sequence alignment was performed using the Geneious prime application and annotations of K96 genes were individually inspected to identify inactivating mutations.

### Statistical analysis

Survival of bacteria in blood and serum, β-galactosidase, and *Galleria mellonella* infection assay data were analyzed using GraphPad Prism 5.0. The statistical tests performed as well as the significance values are indicated in the individual figure legends.

## Acknowledgements

This work was funded by NIAID R15AI159847-01A1 to DE. The funders had no role in study design, data collection and interpretation, or the decision to submit the work for publication.

## References

1. Collaborators GBDAR. Global mortality associated with 33 bacterial pathogens in 2019: a systematic analysis for the Global Burden of Disease Study 2019. Lancet. 2022;400(10369):2221–48. Epub 20221121. doi: 10.1016/S0140-6736(22)02185-7. PubMed PMID: 36423648; PubMed Central PMCID: PMCPMC9763654.

2. Klein RD, Hultgren SJ. Urinary tract infections: microbial pathogenesis, host-pathogen interactions and new treatment strategies. Nat Rev Microbiol. 2020;18(4):211–26. Epub 20200218. doi: 10.1038/s41579-020-0324-0. PubMed PMID: 32071440; PubMed Central PMCID: PMCPMC7942789.

3. Belanger L, Garenaux A, Harel J, Boulianne M, Nadeau E, Dozois CM. Escherichia coli from animal reservoirs as a potential source of human extraintestinal pathogenic E. coli. FEMS Immunol Med Microbiol. 2011;62(1):1–10. Epub 20110324. doi: 10.1111/j.1574-695X.2011.00797.x. PubMed PMID: 21362060.

4. Li G, Cai W, Hussein A, Wannemuehler YM, Logue CM, Nolan LK. Proteome response of an extraintestinal pathogenic Escherichia coli strain with zoonotic potential to human and chicken sera. J Proteomics. 2012;75(15):4853–62. doi: 10.1016/j.jprot.2012.05.044. PubMed PMID: 22677113.

5. Hammad AM, Gonzalez-Escalona N, El Tahan A, Abbas NH, Koenig SSK, Allue-Guardia A, et al. Pathogenome comparison and global phylogeny of Escherichia coli ST1485 strains. Scientific reports. 2022;12(1):18495. Epub 20221102. doi: 10.1038/s41598-022-20342-0. PubMed PMID: 36323726; PubMed Central PMCID: PMCPMC9630279.

6. Jorgensen SL, Stegger M, Kudirkiene E, Lilje B, Poulsen LL, Ronco T, et al. Diversity and Population Overlap between Avian and Human Escherichia coli Belonging to Sequence Type 95. mSphere. 2019;4(1). Epub 20190116. doi: 10.1128/mSphere.00333-18. PubMed PMID: 30651401; PubMed Central PMCID: PMCPMC6336079.

7. Pitout JD. Extraintestinal pathogenic Escherichia coli: an update on antimicrobial resistance, laboratory diagnosis and treatment. Expert review of anti-infective therapy. 2012;10(10):1165–76. doi: 10.1586/eri.12.110. PubMed PMID: 23199402.

8. Pitout JD, Laupland KB. Extended-spectrum beta-lactamase-producing Enterobacteriaceae: an emerging public-health concern. Lancet Infect Dis. 2008;8(3):159–66. Epub 2008/02/23. doi: 10.1016/S1473-3099(08)70041-0. PubMed PMID: 18291338.

9. Antimicrobial Resistance C. Global burden of bacterial antimicrobial resistance in 2019: a systematic analysis. Lancet. 2022;399(10325):629–55. Epub 20220119. doi: 10.1016/S0140-6736(21)02724-0. PubMed PMID: 35065702; PubMed Central PMCID: PMCPMC8841637.

10. Geurtsen J, de Been M, Weerdenburg E, Zomer A, McNally A, Poolman J. Genomics and pathotypes of the many faces of Escherichia coli. FEMS Microbiol Rev. 2022;46(6). doi: 10.1093/femsre/fuac031. PubMed PMID: 35749579; PubMed Central PMCID: PMCPMC9629502.

11. Jann K, Jann B. Capsules of Escherichia coli, expression and biological significance. Can J Microbiol. 1992;38(7):705–10. doi: 10.1139/m92-116. PubMed PMID: 1393836.

12. Whitfield C. Biosynthesis and assembly of capsular polysaccharides in Escherichia coli. Annu Rev Biochem. 2006;75:39–68. doi: 10.1146/annurev.biochem.75.103004.142545. PubMed PMID: 16756484.

13. Sande C, Whitfield C. Capsules and Extracellular Polysaccharides in Escherichia coli and Salmonella. EcoSal Plus. 2021;9(2):eESP00332020. Epub 20211201. doi: 10.1128/ecosalplus.ESP-0033-2020. PubMed PMID: 34910576.

14. Willis LM, Whitfield C. Structure, biosynthesis, and function of bacterial capsular polysaccharides synthesized by ABC transporter-dependent pathways. Carbohydr Res. 2013;378:35–44. doi: 10.1016/j.carres.2013.05.007. PubMed PMID: 23746650.

15. Buckles EL, Wang X, Lane MC, Lockatell CV, Johnson DE, Rasko DA, et al. Role of the K2 capsule in Escherichia coli urinary tract infection and serum resistance. J Infect Dis. 2009;199(11):1689–97. doi: 10.1086/598524. PubMed PMID: 19432551; PubMed Central PMCID: PMCPMC3872369.

16. Johnson JR, Russo TA. Molecular Epidemiology of Extraintestinal Pathogenic Escherichia coli. EcoSal Plus. 2004;1(1). Epub 2004/12/01. doi: 10.1128/ecosalplus.8.6.1.4. PubMed PMID: 26443356.

17. Anderson GG, Goller CC, Justice S, Hultgren SJ, Seed PC. Polysaccharide capsule and sialic acid-mediated regulation promote biofilm-like intracellular bacterial communities during cystitis. Infect Immun. 2010;78(3):963–75. Epub 2010/01/21. doi: 10.1128/IAI.00925-09. PubMed PMID: 20086090; PubMed Central PMCID: PMCPMC2825929.

18. Thomassin JL, Lee MJ, Brannon JR, Sheppard DC, Gruenheid S, Le Moual H. Both group 4 capsule and lipopolysaccharide O-antigen contribute to enteropathogenic Escherichia coli resistance to human alpha-defensin 5. PLoS ONE. 2013;8(12):e82475. Epub 20131204. doi: 10.1371/journal.pone.0082475. PubMed PMID: 24324796; PubMed Central PMCID: PMCPMC3853201.

19. Sarkar S, Ulett GC, Totsika M, Phan MD, Schembri MA. Role of capsule and O antigen in the virulence of uropathogenic Escherichia coli. PLoS ONE. 2014;9(4):e94786. doi: 10.1371/journal.pone.0094786. PubMed PMID: 24722484; PubMed Central PMCID: PMCPMC3983267.

20. Schneider G, Dobrindt U, Bruggemann H, Nagy G, Janke B, Blum-Oehler G, et al. The pathogenicity island-associated K15 capsule determinant exhibits a novel genetic structure and correlates with virulence in uropathogenic Escherichia coli strain 536. Infect Immun. 2004;72(10):5993–6001. doi: 10.1128/IAI.72.10.5993-6001.2004. PubMed PMID: 15385503; PubMed Central PMCID: PMCPMC517556.

21. Russo TA, Liang Y, Cross AS. The presence of K54 capsular polysaccharide increases the pathogenicity of Escherichia coli in vivo. J Infect Dis. 1994;169(1):112–8. Epub 1994/01/01. doi: 10.1093/infdis/169.1.112. PubMed PMID: 8277173.

22. Bahrani-Mougeot FK, Buckles EL, Lockatell CV, Hebel JR, Johnson DE, Tang CM, et al. Type 1 fimbriae and extracellular polysaccharides are preeminent uropathogenic Escherichia coli virulence determinants in the murine urinary tract. Mol Microbiol. 2002;45(4):1079–93. Epub 2002/08/16. doi: 10.1046/j.1365-2958.2002.03078.x. PubMed PMID: 12180926.

23. An H, Qian C, Huang Y, Li J, Tian X, Feng J, et al. Functional vulnerability of liver macrophages to capsules defines virulence of blood-borne bacteria. The Journal of experimental medicine. 2022;219(4). Epub 20220308. doi: 10.1084/jem.20212032. PubMed PMID: 35258552; PubMed Central PMCID: PMCPMC8908791.

24. Aldawood E, Roberts IS. Regulation of Escherichia coli Group 2 Capsule Gene Expression: A Mini Review and Update. Front Microbiol. 2022;13:858767. Epub 20220310. doi: 10.3389/fmicb.2022.858767. PubMed PMID: 35359738; PubMed Central PMCID: PMCPMC8960920.

25. Goh KGK, Phan MD, Forde BM, Chong TM, Yin WF, Chan KG, et al. Genome-Wide Discovery of Genes Required for Capsule Production by Uropathogenic Escherichia coli. mBio. 2017;8(5). Epub 2017/10/27. doi: 10.1128/mBio.01558-17. PubMed PMID: 29066548; PubMed Central PMCID: PMCPMC5654933.

26. Corbett D, Bennett HJ, Askar H, Green J, Roberts IS. SlyA and H-NS regulate transcription of the Escherichia coli K5 capsule gene cluster, and expression of slyA in Escherichia coli is temperature-dependent, positively autoregulated, and independent of H-NS. J Biol Chem. 2007;282(46):33326–35. Epub 20070907. doi: 10.1074/jbc.M703465200. PubMed PMID: 17827501.

27. Stevens MP, Clarke BR, Roberts IS. Regulation of the Escherichia coli K5 capsule gene cluster by transcription antitermination. Mol Microbiol. 1997;24(5):1001–12. doi: 10.1046/j.1365-2958.1997.4241780.x. PubMed PMID: 9220007.

28. Olson MA, Grimsrud A, Richards AC, Mulvey MA, Wilson E, Erickson DL. Bile Salts Regulate Zinc Uptake and Capsule Synthesis in a Mastitis-Associated Extraintestinal Pathogenic Escherichia coli Strain. Infect Immun. 2021;89(10):e0035721. Epub 20210706. doi: 10.1128/IAI.00357-21. PubMed PMID: 34228495; PubMed Central PMCID: PMCPMC8445175.

29. Riley LW. Pandemic lineages of extraintestinal pathogenic Escherichia coli. Clin Microbiol Infect. 2014;20(5):380–90. doi: 10.1111/1469-0691.12646. PubMed PMID: 24766445.

30. Olson MA, Siebach TW, Griffitts JS, Wilson E, Erickson DL. Genome-Wide Identification of Fitness Factors in Mastitis-Associated Escherichia coli. Appl Environ Microbiol. 2018;84(2):e02190–17. Epub 2017/11/05. doi: 10.1128/AEM.02190-17. PubMed PMID: 29101196; PubMed Central PMCID: PMCPMC5752858.

31. Yang S, Xi D, Jing F, Kong D, Wu J, Feng L, et al. Genetic diversity of K-antigen gene clusters of Escherichia coli and their molecular typing using a suspension array. Can J Microbiol. 2018;64(4):231–41. Epub 2018/01/23. doi: 10.1139/cjm-2017-0620. PubMed PMID: 29357266.

32. Jann B, Kochanowski H, Jann K. Structure of the capsular K96 polysaccharide (K96 antigen) from Escherichia coli O77:K96:H– and comparison with the capsular K54 polysaccharide (K54 antigen) from Escherichia coli O6:K54:H10. Carbohydr Res. 1994;253:323–7. Epub 1994/02/03. doi: 10.1016/0008-6215(94)80080-4. PubMed PMID: 8156556.

33. Hofmann P, Jann B, Jann K. Structure of the Amino Acid-Containing Capsular Polysaccharide (K54-Antigen) from Escherichia-Coli O6-K54-H10. Carbohydr Res. 1985;139(Jun):261–71. PubMed PMID: WOS:A1985ALS7000025.

34. Russo T, Brown JJ, Jodush ST, Johnson JR. The O4 specific antigen moiety of lipopolysaccharide but not the K54 group 2 capsule is important for urovirulence of an extraintestinal isolate of Escherichia coli. Infect Immun. 1996;64(6):2343–8. Epub 1996/06/01. PubMed PMID: 8675348; PubMed Central PMCID: PMCPMC174077.

35. Gladstone RA, Pesonen M, Pöntinen AK, Mäklin T, MacAlasdair N, Thorpe H, et al. Group 2 and 3 ABC-transporter dependant capsular K-loci contribute significantly to variation in the invasive potential of Escherichia coli. medRxiv. 2024:2024.11.22.24317484.

36. Kim KS, Itabashi H, Gemski P, Sadoff J, Warren RL, Cross AS. The K1 capsule is the critical determinant in the development of Escherichia coli meningitis in the rat. J Clin Invest. 1992;90(3):897–905. Epub 1992/09/01. doi: 10.1172/JCI115965. PubMed PMID: 1326000; PubMed Central PMCID: PMCPMC329944.

37. Leying H, Suerbaum S, Kroll HP, Stahl D, Opferkuch W. The capsular polysaccharide is a major determinant of serum resistance in K-1-positive blood culture isolates of Escherichia coli. Infect Immun. 1990;58(1):222–7. Epub 1990/01/01. PubMed PMID: 2403532; PubMed Central PMCID: PMCPMC258433.

38. Bjorksten B, Bortolussi R, Gothefors L, Quie PG. Interaction of E. coli strains with human serum: lack of relationship to K1 antigen. J Pediatr. 1976;89(6):892–7. Epub 1976/12/01. doi: 10.1016/s0022-3476(76)80592-6. PubMed PMID: 792409.

39. Russo TA, Moffitt MC, Hammer CH, Frank MM. TnphoA-mediated disruption of K54 capsular polysaccharide genes in Escherichia coli confers serum sensitivity. Infect Immun. 1993;61(8):3578–82. Epub 1993/08/01. PubMed PMID: 8392976; PubMed Central PMCID: PMCPMC281046.

40. Wetsel RA, Fleischer DT, Haviland DL. Deficiency of the murine fifth complement component (C5). A 2-base pair gene deletion in a 5’-exon. J Biol Chem. 1990;265(5):2435–40. PubMed PMID: 2303408.

41. Dorman MJ, Feltwell T, Goulding DA, Parkhill J, Short FL. The Capsule Regulatory Network of Klebsiella pneumoniae Defined by density-TraDISort. mBio. 2018;9(6). Epub 20181120. doi: 10.1128/mBio.01863-18. PubMed PMID: 30459193; PubMed Central PMCID: PMCPMC6247091.

42. Hammerschmidt S, Muller A, Sillmann H, Muhlenhoff M, Borrow R, Fox A, et al. Capsule phase variation in Neisseria meningitidis serogroup B by slipped-strand mispairing in the polysialyltransferase gene (siaD): correlation with bacterial invasion and the outbreak of meningococcal disease. Mol Microbiol. 1996;20(6):1211–20. doi: 10.1111/j.1365-2958.1996.tb02641.x. PubMed PMID: 8809773.

43. Waite RD, Penfold DW, Struthers JK, Dowson CG. Spontaneous sequence duplications within capsule genes cap8E and tts control phase variation in Streptococcus pneumoniae serotypes 8 and 37. Microbiology (Reading). 2003;149(Pt 2):497–504. doi: 10.1099/mic.0.26011-0. PubMed PMID: 12624211.

44. Bailey MJ, Hughes C, Koronakis V. RfaH and the *ops* element, components of a novel system controlling bacterial transcription elongation. Mol Microbiol. 1997;26(5):845–51. doi: 10.1046/j.1365-2958.1997.6432014.x. PubMed PMID: 9426123.

45. Lippolis JD, Holman DB, Brunelle BW, Thacker TC, Bearson BL, Reinhardt TA, et al. Genomic and Transcriptomic Analysis of Escherichia coli Strains Associated with Persistent and Transient Bovine Mastitis and the Role of Colanic Acid. Infect Immun. 2018;86(1). Epub 2017/10/25. doi: 10.1128/IAI.00566-17. PubMed PMID: 29061709; PubMed Central PMCID: PMCPMC5736815.

46. Germon P, Foucras G, Smith DGE, Rainard P. Invited review: Mastitis Escherichia coli strains-Mastitis-associated or mammo-pathogenic? J Dairy Sci. 2025;108(5):4485–507. Epub 20250324. doi: 10.3168/jds.2024-26109. PubMed PMID: 40139360.

47. Johnson JR, Russo TA. Molecular Epidemiology of Extraintestinal Pathogenic Escherichia coli. EcoSal Plus. 2018;8(1). Epub 2018/04/19. doi: 10.1128/ecosalplus.ESP-0004-2017. PubMed PMID: 29667573.

48. Hong Y, Qin J, Forga XB, Totsika M. Extensive diversity in Escherichia coli group 3 capsules is driven by recombination and plasmid transfer from multiple species. Microbiology Spectrum. 2023;11(4):e01432–23.

49. Serruto D, Rappuoli R, Scarselli M, Gros P, van Strijp JA. Molecular mechanisms of complement evasion: learning from staphylococci and meningococci. Nat Rev Microbiol. 2010;8(6):393–9. doi: 10.1038/nrmicro2366. PubMed PMID: 20467445.

50. Meri S, Pangburn MK. Discrimination between activators and nonactivators of the alternative pathway of complement: regulation via a sialic acid/polyanion binding site on factor H. Proc Natl Acad Sci U S A. 1990;87(10):3982–6. doi: 10.1073/pnas.87.10.3982. PubMed PMID: 1692629; PubMed Central PMCID: PMCPMC54028.

51. McCarthy AJ, Stabler RA, Taylor PW. Genome-Wide Identification by Transposon Insertion Sequencing of Escherichia coli K1 Genes Essential for In Vitro Growth, Gastrointestinal Colonizing Capacity, and Survival in Serum. J Bacteriol. 2018;200(7). Epub 2018/01/18. doi: 10.1128/JB.00698-17. PubMed PMID: 29339415; PubMed Central PMCID: PMCPMC5847654.

52. Salamon H, Nissim-Eliraz E, Ardronai O, Nissan I, Shpigel NY. The role of O-polysaccharide chain and complement resistance of Escherichia coli in mammary virulence. Veterinary research. 2020;51(1):77. Epub 20200615. doi: 10.1186/s13567-020-00804-x. PubMed PMID: 32539761; PubMed Central PMCID: PMCPMC7294653.

53. Unverdorben LV, Pirani A, Gontjes K, Moricz B, Holmes CL, Snitkin ES, et al. Klebsiella pneumoniae evolution in the gut leads to spontaneous capsule loss and decreased virulence potential. mBio. 2025;16(5):e0236224. Epub 20250331. doi: 10.1128/mbio.02362-24. PubMed PMID: 40162782; PubMed Central PMCID: PMCPMC12077207.

54. Wei DW, Song Y, Li Y, Zhang G, Chen Q, Wu L, et al. Insertion sequences accelerate genomic convergence of multidrug resistance and hypervirulence in Klebsiella pneumoniae via capsular phase variation. Genome Med. 2025;17(1):45. Epub 20250506. doi: 10.1186/s13073-025-01474-0. PubMed PMID: 40329368; PubMed Central PMCID: PMCPMC12057282.

55. Almeida RA, Luther DA, Kumar SJ, Calvinho LF, Bronze MS, Oliver SP. Adherence of Streptococcus uberis to bovine mammary epithelial cells and to extracellular matrix proteins. Zentralbl Veterinarmed B. 1996;43(7):385–92. doi: 10.1111/j.1439-0450.1996.tb00330.x. PubMed PMID: 8885703.

56. Fujisaki S, Hara H, Nishimura Y, Horiuchi K, Nishino T. Cloning and nucleotide sequence of the ispA gene responsible for farnesyl diphosphate synthase activity in Escherichia coli. J Biochem. 1990;108(6):995–1000. doi: 10.1093/oxfordjournals.jbchem.a123327. PubMed PMID: 2089044.

57. Workman SD, Strynadka NCJ. A Slippery Scaffold: Synthesis and Recycling of the Bacterial Cell Wall Carrier Lipid. J Mol Biol. 2020;432(18):4964–82. Epub 20200329. doi: 10.1016/j.jmb.2020.03.025. PubMed PMID: 32234311.

58. Seo SW, Kim D, Szubin R, Palsson BO. Genome-wide Reconstruction of OxyR and SoxRS Transcriptional Regulatory Networks under Oxidative Stress in Escherichia coli K-12 MG1655. Cell Rep. 2015;12(8):1289–99. Epub 20150813. doi: 10.1016/j.celrep.2015.07.043. PubMed PMID: 26279566.

59. Pickard D, Kingsley RA, Hale C, Turner K, Sivaraman K, Wetter M, et al. A genomewide mutagenesis screen identifies multiple genes contributing to Vi capsular expression in Salmonella enterica serovar Typhi. J Bacteriol. 2013;195(6):1320–6. Epub 20130111. doi: 10.1128/JB.01632-12. PubMed PMID: 23316043; PubMed Central PMCID: PMCPMC3592008.

60. Lee GY, Song J. Single missense mutations in Vi capsule synthesis genes confer hypervirulence to Salmonella Typhi. Nat Commun. 2024;15(1):5258. Epub 20240619. doi: 10.1038/s41467-024-49590-6. PubMed PMID: 38898034; PubMed Central PMCID: PMCPMC11187135.

61. Hancock SJ, Lo AW, Ve T, Day CJ, Tan L, Mendez AA, et al. Ucl fimbriae regulation and glycan receptor specificity contribute to gut colonisation by extra-intestinal pathogenic Escherichia coli. PLoS Pathog. 2022;18(6):e1010582.

62. Elazar S, Gonen E, Livneh-Kol A, Rosenshine I, Shpigel NY. Essential role of neutrophils but not mammary alveolar macrophages in a murine model of acute Escherichia coli mastitis. Veterinary research. 2010;41(4):53. doi: 10.1051/vetres/2010025. PubMed PMID: 20416261; PubMed Central PMCID: PMCPMC2881416.

63. Condron C, Toomey D, Casey RG, Shaffii M, Creagh T, Bouchier-Hayes D. Neutrophil bactericidal function is defective in patients with recurrent urinary tract infections. Urol Res. 2003;31(5):329–34. Epub 2003/10/24. doi: 10.1007/s00240-003-0344-z. PubMed PMID: 14574538.

64. Zhao C, Hartke A, La Sorda M, Posteraro B, Laplace JM, Auffray Y, et al. Role of methionine sulfoxide reductases A and B of Enterococcus faecalis in oxidative stress and virulence. Infect Immun. 2010;78(9):3889–97. Epub 2010/06/23. doi: IAI.00165-10 [pii] 10.1128/IAI.00165-10. PubMed PMID: 20566694; PubMed Central PMCID: PMC2937430.

65. Erickson DL, Russell CW, Johnson KL, Hileman T, Stewart RM. PhoP and OxyR transcriptional regulators contribute to Yersinia pestis virulence and survival within Galleria mellonella. Microb Pathog. 2011. Epub 2011/10/04. doi: 10.1016/j.micpath.2011.08.008. PubMed PMID: 21964409.

66. Thompson CD, Khan MF, Crosby LRG, Holcomb SG, Vidal AGJ, Vidal JE, et al. AliC and AliD of nonencapsulated Streptococcus pneumoniae enhance virulence in a Galleria mellonella model of infection by contributing to reactive oxygen species resistance. Front Cell Infect Microbiol. 2025;15:1583375. Epub 20250611. doi: 10.3389/fcimb.2025.1583375. PubMed PMID: 40568701; PubMed Central PMCID: PMCPMC12187840.

67. Erickson DL, Lew CS, Kartchner B, Porter NT, McDaniel SW, Jones NM, et al. Lipopolysaccharide Biosynthesis Genes of Yersinia pseudotuberculosis Promote Resistance to Antimicrobial Chemokines. PLoS ONE. 2016;11(6):e0157092. doi: 10.1371/journal.pone.0157092. PubMed PMID: 27275606; PubMed Central PMCID: PMCPMC4898787.

68. Olson MA, Cullimore C, Hutchison WD, Grimsrud A, Nobrega D, De Buck J, et al. Genes associated with fitness and disease severity in the pan-genome of mastitis-associated Escherichia coli. Front Microbiol. 2024;15:1452007. Epub 20240829. doi: 10.3389/fmicb.2024.1452007. PubMed PMID: 39268542; PubMed Central PMCID: PMCPMC11390585.

69. Dobin A, Davis CA, Schlesinger F, Drenkow J, Zaleski C, Jha S, et al. STAR: ultrafast universal RNA-seq aligner. Bioinformatics. 2013;29(1):15–21. Epub 20121025. doi: 10.1093/bioinformatics/bts635. PubMed PMID: 23104886; PubMed Central PMCID: PMCPMC3530905.

70. Cianfanelli FR, Cunrath O, Bumann D. Efficient dual-negative selection for bacterial genome editing. BMC Microbiol. 2020;20(1):129. Epub 20200524. doi: 10.1186/s12866-020-01819-2. PubMed PMID: 32448155; PubMed Central PMCID: PMCPMC7245781.

71. Parveen N, Durrant S, Olson MA, Shakespear EP, Jones TR, Wilson E, et al. Opposing roles for iron transport systems in gallium tolerance in extraintestinal pathogenic Escherichia coli. bioRxiv. 2025:2025.07.01.662525. doi: 10.1101/2025.07.01.662525.

72. Zhang X, Bremer H. Control of the Escherichia coli rrnB P1 promoter strength by ppGpp. J Biol Chem. 1995;270(19):11181–9. doi: 10.1074/jbc.270.19.11181. PubMed PMID: 7538113.

